# Survival of the frequent at finite population size and mutation rate: bridging the gap between quasispecies and monomorphic regimes with a simple model

**DOI:** 10.1101/375147

**Authors:** Bhavin S. Khatri

## Abstract

In recent years, there has been increased attention on the non-trivial role that genotype-phenotype maps play in the course of evolution, where natural selection acts on phenotypes, but variation arises at the level of mutations. Understanding such mappings is arguably the next missing piece in a fully predictive theory of evolution. Although there are theoretical descriptions of such mappings for the monomorphic (*Nμ* ≪ 1) and deterministic or very strong mutation (*Nμ* ⋙ 1) limit, given by developments of Iwasa’s free fitness and quasispecies theories, respectively, there is no general description for the intermediate regime where *Nμ* ~ 1. In this paper, we address this by transforming Wright’s well-known stationary distribution of genotypes under selection and mutation to give the probability distribution of phenotypes, assuming a general genotype-phenotype map. The resultant distribution shows that the degeneracies of each phenotype appear by weighting the mutation term; this gives rise to a bias towards phenotypes of larger degeneracy analogous to quasispecies theory, but at finite population size. On the other hand we show that as population size is decreased, again phenotypes of higher degeneracy are favoured, which is a finite mutation description of the effect of sequence entropy in the monomorphic limit. We also for the first time (to the author’s knowledge) provide an explicit derivation of Wright’s stationary distribution of the frequencies of multiple alleles.

## INTRODUCTION

The past 100 years has seen our understanding of the mechanisms of evolution develop, from its initial mathematical foundations due initially to Wright and Fisher, and later Kimura, which encompass a description of the interplay between selection, mutation and drift, to the current day with descriptions of multi-locus evolution with recombination, linkage and epistasis^1^. However, as powerful as these studies are, they lack a crucial missing ingredient in our understanding of evolution, which is the role of genotype-phenotype maps, where selection acts on phenotypes, but underlying variation arises at the genetic level. In general, such mappings will be very complex, but a common theme is that because these mappings will often be many-to-one, some phenotypes will have more genotypes associated with them than others. In the weak mutation or monomorphic limit, where the population-scaled mutation rate is small (*Nμ* ≪ 1), there is a complete theory that predicts the equilibrium distribution of phenotypes in the monomorphic limit^2^, which naturally leads from Iwasa’s definition of *free fitness*^3^ (subsequently rediscovered by Sella & Hirsh^4^). A general prediction of these theories is that at small population sizes, as genetic drift dominates favouring phenotypes with larger sequence entropy, which is the log degeneracy of the phenotype. In the opposite regime, with infinitely large population sizes there is Eigen’s quasispecies theory, which are deterministic sets of equations describing the growth and mutation of many genotypes^5^; one of its predictions is that at sufficiently large mutation rates regions of locally high robustness in genotype space are favoured^6,7^. Translating to phenotype space, it is trivial to see that phenotypes that have higher (average) local robustness will be favoured over those with lower robustness. In both the monomorphic and quasispecies regimes, we see analogous effects related to non-optimal degenerate (robust) phenotypes being favoured at small population sizes (large mutation rates).

In this paper, we present a theory that straddles both these regimes to calculate the equilibrium distribution of phenotype frequencies at arbitrary and finite population sizes and mutation rates. This is done by transforming Wright’s equilibrium (stationary) distribution of multiples alleles to the space of phenotypes assuming a simple many-to-one genotype to phenotype map. We do this making the simplifying assumption that all genotypes are connected, which means detailed balance is obeyed and Wright’s stationary distribution is valid. The result is a distribution of the same form as the distribution of the frequencies of genotypes, but where the mutation term for each phenotype is weighted by the degeneracy of that phenotype; this shows that phenotypes of high degeneracy will tend to be favoured as on average there are more mutational paths into them. We show explicitly in the case of two phenotypes a phase-transition in frequency to the more degenerate phenotype as population size is reduced and/or mutation rate increased. This theory represent a finite population sized description of the analogous phenomenon to survival of the flattest of quasispecies theory and a strong mutation description of survival of the frequent found in the monomorphic weak mutation regime.

## TRANSFORMING DISTRIBUTION FROM GENOTYPE TO PHENOTYPE

We assume a genotype space denoted by a vector ***g***, where *g_i_* = {*σ_k_*}, where *σ_k_* represents possible symbols at each site *i* of the genome, from an alphabet of size 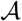. We assume each genotype has fitness *f_**g**_*. The rate of mutations into any given state is then assumed the same as any other state, 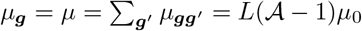. In this case, the equilibrium distribution of the frequency of genotypes, *x_**g**_*:

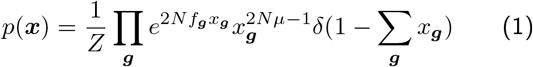

which we derive in the Appendix, as to the author’s knowledge this has not been done explicitly in the literature^8^. Here *N* is the effective population size and the vector ***x*** is a vector of the frequency of genotypes. Given a genotype-phenotype map *ξ* = Ξ(***g***), which we assume is many to one, we want to recast this distribution in terms of the probability distribution of the frequency of *n* phenotypes *z_ξ_* It is clear that the distribution of phenotype frequencies should be of the same form as Eqn. 1, as changing variables to the space of phenotypes does not change the underlying population genetic problem, just the number of alleles to *n*; however, although the fitness of each phenotype is clear, what the effective mutation rate should be is not so clear. To evaluate the distribution explicitly we have:

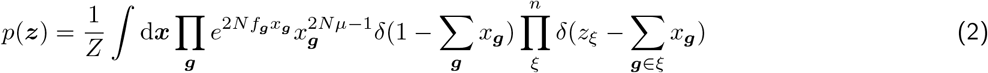

∑_***g***_ *x_**g**_* = ∑_*ξ*_ *z_ξ_* and so the first delta function constraint enforces ∑_*ξ*_ *z_ξ_* = 1. The product over genotypes can be decomposed into a product over phenotypes and a product over genotypes which map to the same phenotype, where these genotypes have the same fitness, by definition, *f_ξ_*:

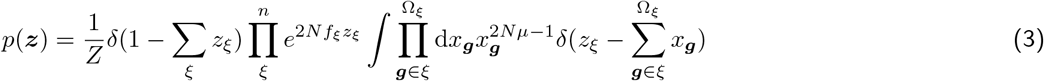

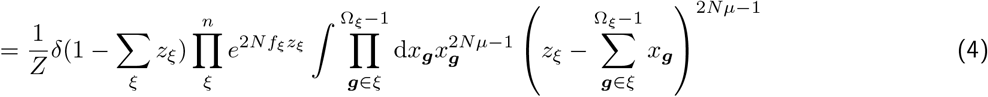

Integrating over the frequency of a single genotype of each phenotype, we are left to evaluate a multidimensional integral over the remaining genotypes, which are coupled due to the constraint that the sum over their frequencies should be *z_ξ_*. To perform the integral we modify the transformation in^9^, which transforms the unit simplex to the unit cube; here we will transform the simplex over all genotypes that belong to a given phenotype constrained to sum to frequency *z_ξ_* to the unit cube over transformed genotype frequencies *u_i_* (switching to a linear index *i* corresponding to the *i^th^* genotype ***g**_i_*):

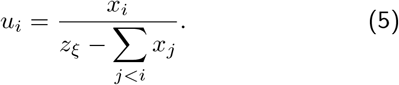

The inverse of this transformation is

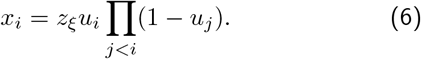

Making the change of variable in the integral, and using the fact that

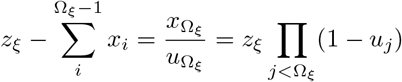

we have:

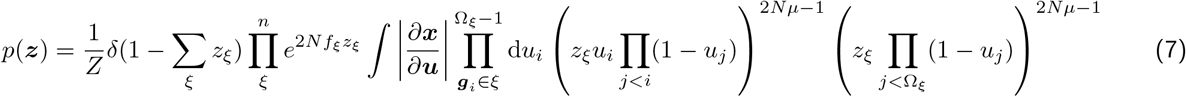

The determinant of the Jacobian can be simply evaluated since |*∂_x_i__/∂_u_j__*| = 0 for *j > i* and so the determinant is the product of the diagonal elements:

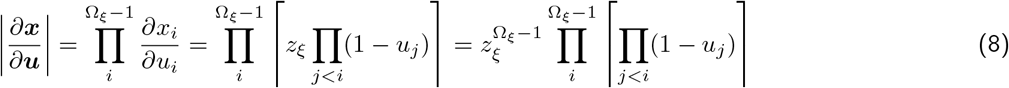

Plugging this into above we get:

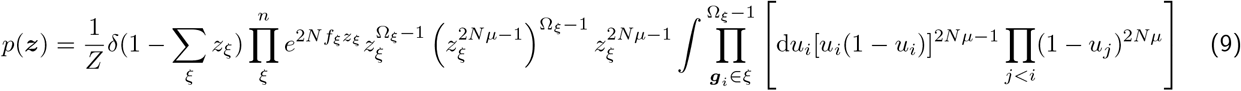

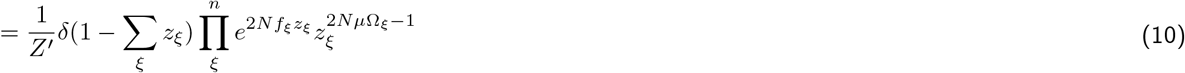

where *Z*′ is the normalisation factor. The key result here is that the degeneracy of phenotypes enhances the effective mutation rate into that phenotype giving a bias to increase the frequency of that phenotype.

## EXAMPLE: TWO PHENOTYPES

Let’s assume there are two phenotypes with log-fitness *f*_1_ and *f*_2_, with selection coefficient *s* = *f*_1_ − *f*_2_, degeneracies Ω_1_, and Ω_2_ and a base-pair mutation rate *μ*. If the frequency of phenotype 1 is denoted *z*, and phenotype 2 1 – *z*, the probability density is given by Eqn.9:

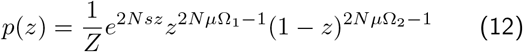

where 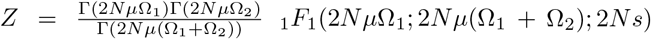. As shown in Fig.1, if we allow phenotype 1 to be advantageous with *s* > 0, there is a shift from phenotype 1 being at high frequency, when the degeneracies are equal (Ω_1_ = Ω_2_) to favouring phenotype 2 (reduction in frequency *z*), when there is a strong bias in the genotype-phenotype map towards phenotype 2 (Ω_2_ ≫ Ω_1_). For large population sizes (a) we are in the strong mutation regime, where 2*N μ*Ω_1_ ≫ 1 and 2*N μ*Ω_2_ ≫ 1, and we see there is a mutation-selection balance in favour of the advantageous phenotype, when degeneracies are equal, and this equilibrium moves to smaller frequencies as the ratio of the degeneracies Ω_2_/Ω_1_ increases; this is an analogous finite *N* description of population delocalisation as found for infinite *N* deterministic quasispecies modelling of populations^6,7^, except here we capture the broad fluctuations around the mutation-selection balance equilibrium. On the other hand when the effect of mutations is weak, 2*N μ*Ω_1_ ≪ 1 and 2*N μ*Ω_2_ ≪ 1, as shown in (b), we see distributions characteristic of the mostly monomorphic composition of the population at any given time, where distributions are condensed at *z* = 0 and *z* = 1; nonetheless we see that when the ratio of the degeneracies (Ω_2_/Ω_1_) becomes large, the distributions shift from a larger density near *z* = 1 to one where a larger density at *z* = 0. This is the correct polymorphic extension of the populations in the monomorphic weak mutation regime, which has recently seen attention using such concepts as free fitness and sequence entropy^2–4,10–12^.

**FIG. 1.**
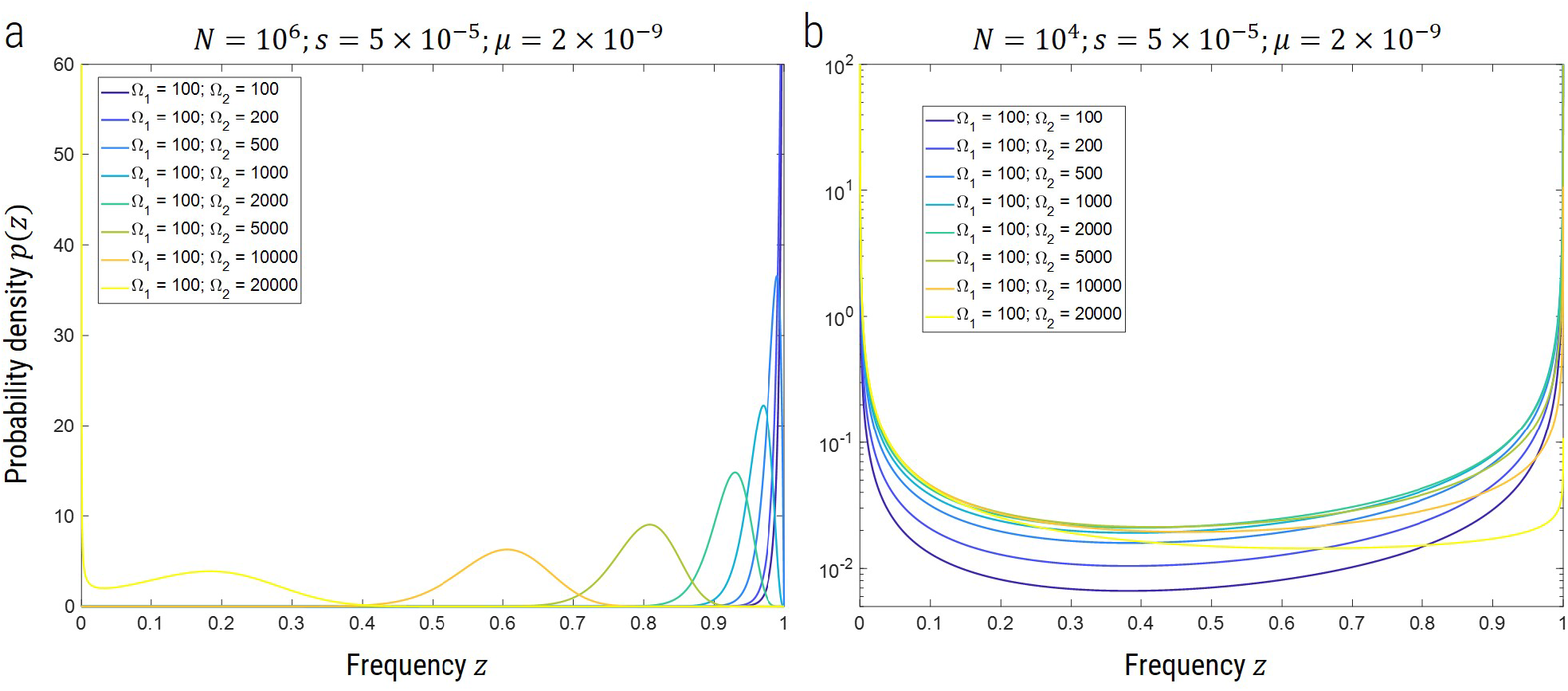
Plot of the phenotypic frequency distribution for different values for *μ* = 2 × 10^−9^, *s* = 5 × 10^−5^ and a) *N* = 10^6^ and b) *N* = 10^4^, for various values of the degeneracy of each phenotype as shown in the legend. N.B. the probability density is plotted on a linear scale in a) and log scale in b) for clarity.

We can also examine the effect of changing the mutation rate or population size in a two phenotype system when there is a large bias in degeneracy. This is most effectively probed by calculating the mean frequency of the phenotype 〈*z*〉 = *∫* d*zzp*(*z*):

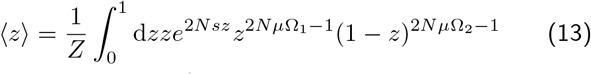

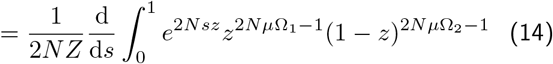

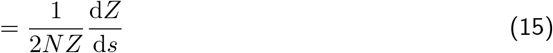

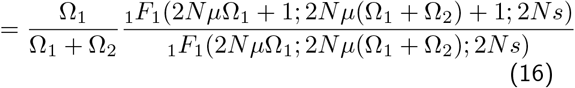

where we have used the fact that 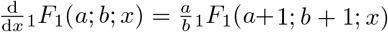. We expect that in the limit that 2*Nμ* → 0 to recover the monomorphic limit^2–4,10^, where the probability of each phenotype is given by 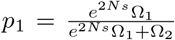 and 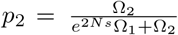, such that 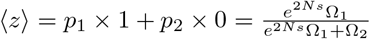. Using the series definition of the confluent hypergeometric function it is straightforward to show that

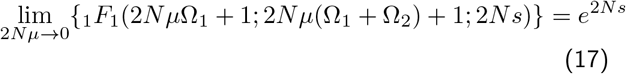

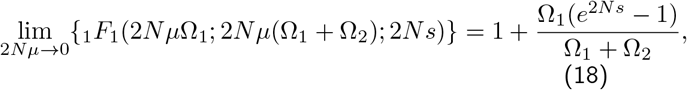

which leads to the following expression for the limit of the mean frequency:

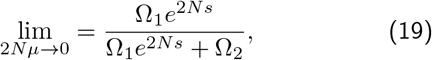

which agrees with the monomorphic expectation.

We can also take the limit 2*N s* → 0, keeping 2*Nμ* finite, which is simply evaluated as lim_*x*→0_{_1_*F*_1_(*a; b; x*)} = 1, giving

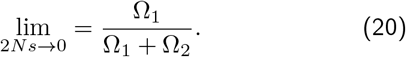

So whether in the monomorphic (2*Nμ* ≪ 1) or polymorphic limit (2*Nμ* > 1), as selection becomes very weak, we get the purely neutral result that the average frequency is determined solely by the relative degeneracies of each phenotype.

In Fig.2, we investigate the mean frequency 〈*z*〉 for Ω_1_ = 100 and Ω_2_ = 10000 and *s* = 5 × 10^−5^, for various values of *N* and *μ*. We see there is a strong delocalisation transition for both *μ* and *N*; as has been essentially described previously for deterministic quasispecies in what is known as the “survival of the flattest”^6,7^, for an increasing mutation rate, we find the less advantageous, but more genotypically degenerate phenotype is favoured. However, here this calculation also shows that concurrently decreasing the effective population size increases progress towards this delocalisation transition, again favouring the more degenerate phenotype.

**FIG. 2.**
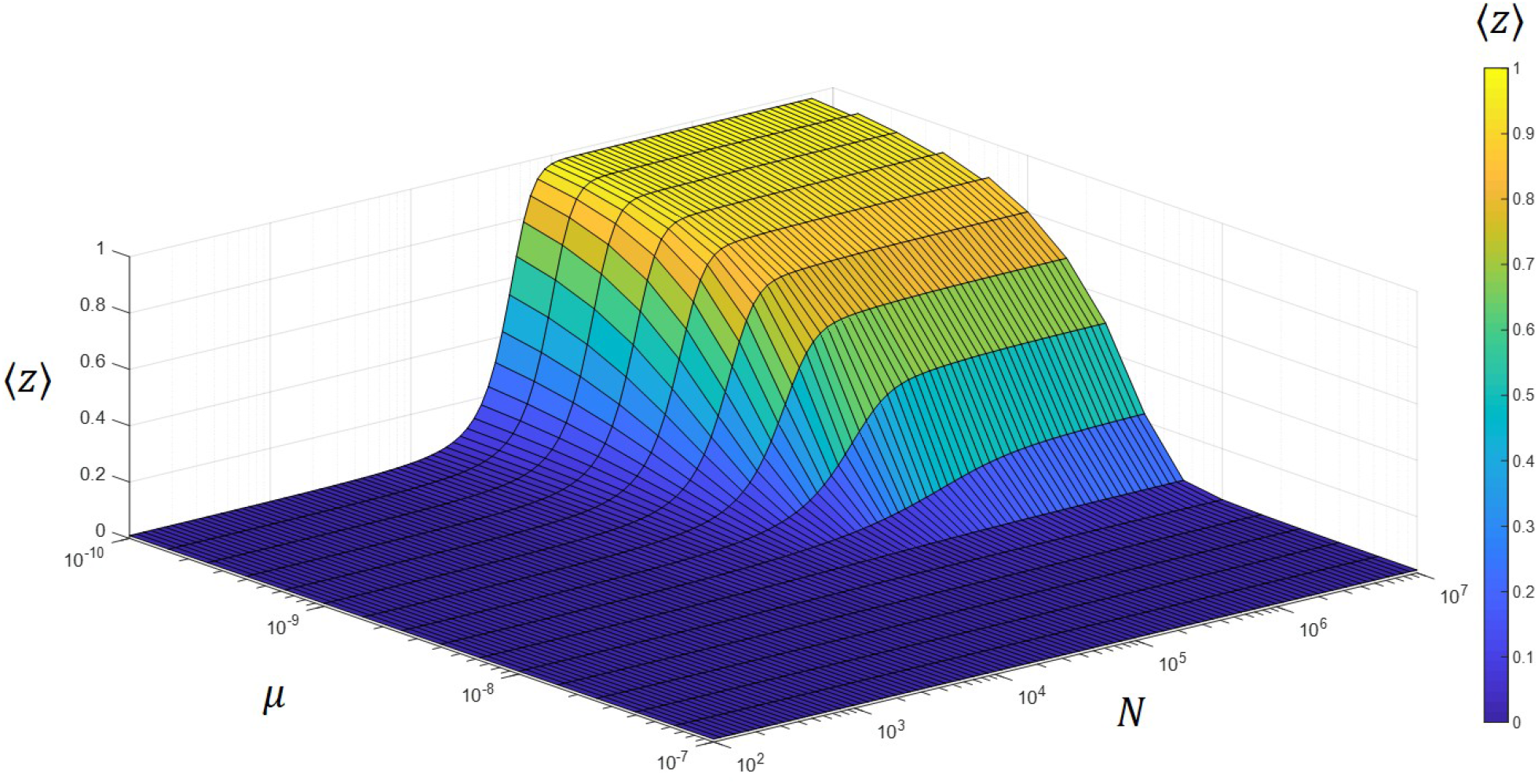
Plot of the mean frequency of phenotype 1, 〈*z*〉, as function of effective population size *N* and mutation rate *μ* for a selection coefficient *s* = 5 × 10^−5^ (favouring phenotype 1) and degeneracies Ω_1_ = 100 and Ω_2_ = 10000.

In Fig.3, we have an equivalent plot for the case when the degeneracies of each phenotype is the same. We see that we have a transition to the neutral state for increasing *μ* or decreasing *N*, giving an equal likelihood of each phenotype, signified by 〈*z*〉 = 1/2.

**FIG. 3.**
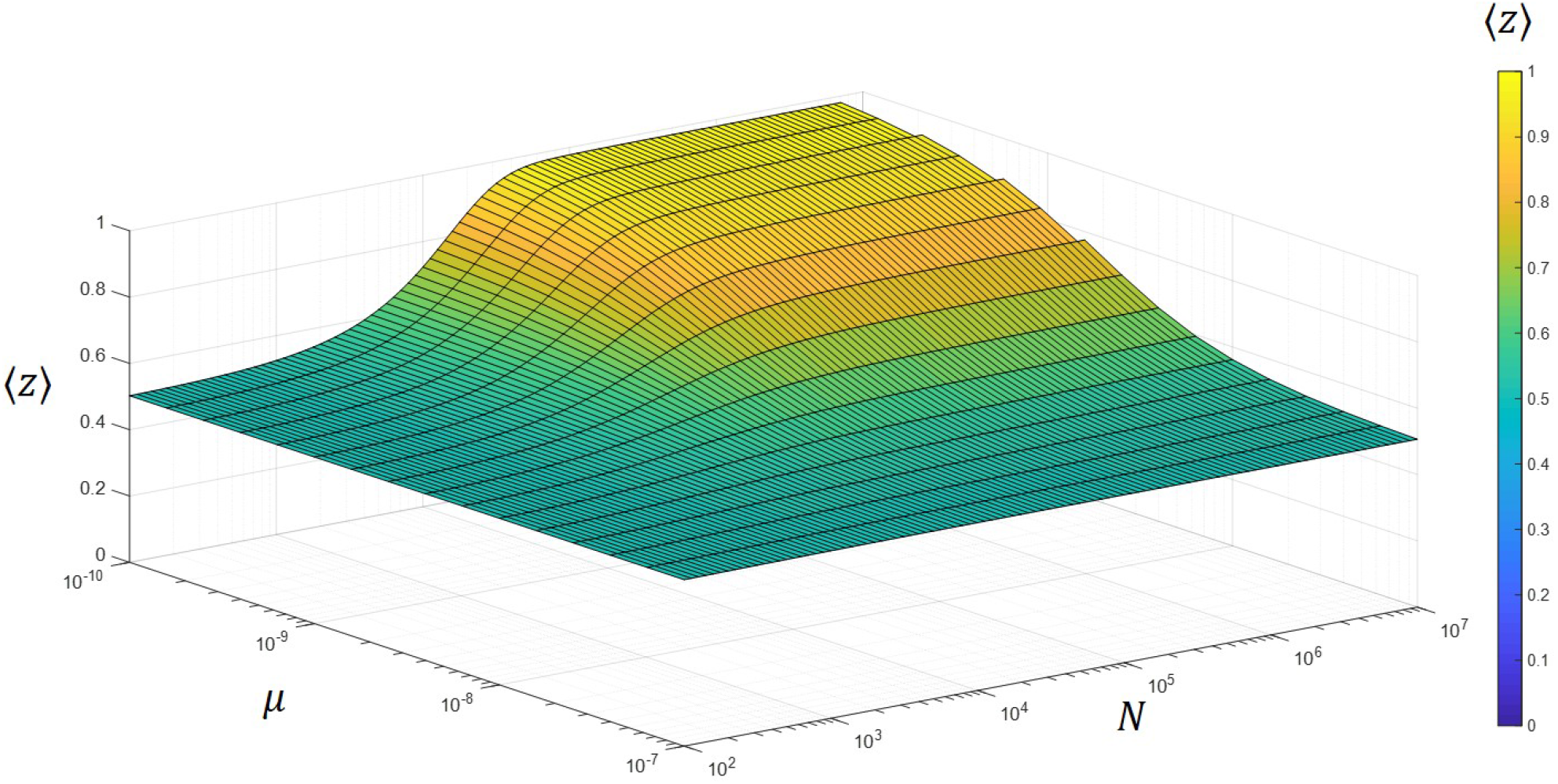
Plot of the mean frequency of phenotype 1, 〈*z*〉, as function of effective population size *N* and mutation rate *μ* for a selection coefficient *s* = 5 × 10^−5^ (favouring phenotype 1) and degeneracies Ω_1_ = 10000 and Ω_2_ = 10000.

## DISCUSSION & CONCLUSIONS

In this paper, we have derived the equilibrium distribution of the frequency of phenotypes, assuming a general many-to-one genotype phenotype map and that the mutation rates between all genotypes is uniform. The result shows that the equilibrium distribution is of the same form as genotypes but the mutation terms are weighted by the degeneracy of each phenotype. This gives rise to a bias towards phenotypes of higher degeneracy as the mutation rate is increased and/or the population size decreased, as we show explicitly for the two phenotype case. This calculation generalises the equilibrium distribution of phenotypes in the monomorphic regime to the polymorphic, describing the analogous effect of the increasing effect of sequence entropy (log degeneracy of phenotypes), as population size is decreased. However, here we note that there is no obvious way to express the equilibrium frequency distribution in terms of an analogous quantity such as sequence entropy.

This calculation is also a finite population size description of the quasispecies limit, which are deterministic infinite population size models^5^. Quasispecies models predict the phenomenon of *survival of the flattest*, which is a delocalisation transition due to higher robustness or smaller ‘local curvature’ being favoured at larger mutation rates^6,7^; here we make no explicit statement about robustness, and we see an analogous phenomenon arises irrespective of how different genotypes of the same phenotype are connected; the difference between this and the quasispecies is that here we have an equilibrium calculation, which assumes a certain ergodicity or accessibility of a representative number of states for each phenotype. Whether this equilibrium is reached on relevant evolutionary timescales is an open question; as discussed in^13^, certain phenotypes may be more likely to arrive as they arise more frequently in the local neighbourhood. On the other hand there are broad reasons to expect the structure of genotype-phenotype maps to be ergodic^14^; in particular, simulations of the genotype-phenotype map for spatial patterning in development studied in^11^ was found to be ergodic, despite simulation times which could not exhaustively search the whole genotype space.

We also present for the first time, to the author’s knowledge, an explicit derivation of Wright’s multi-allele frequency distribution^8^. The derivation makes clear the “curl-free” or “potential” assumption that gives rise to the stationary or equilibrium distribution of allele frequencies. In other words there cannot be any circulating fluxes of probability in the equilibrium state; this arises due to the uniform mutation assumption, and we expect that for arbitrary mutation structure the curl-free assumption will not be satisfied. In particular, a more realistic, but analytically intractable model is one where mutation is between nearest neighbours only; this violates Wright’s assumptions and suggests detailed balance is not obeyed. It is an open question as to which forms of mutation structure can give rise to potential solutions of the form of Wright’s stationary distribution. We believe the results of this paper will likely be robust to such considerations in the limit of large degeneracies of each phenotype, where the product of degeneracy and mutation rate into a phenotype becomes some effective average mutation rate over microscopic mutation rates.

To conclude these results provide a theory to calculate the equilibrium distribution of the frequency of phenotypes for a general genotype-phenotype map with the simplifying assumption of uniform mutation between genotypes. As such it provides a framework to understand the bridge between the weak mutation, monomorphic, regime and infinite population size, deterministic, quasispecies regime, both showing that non-optimal selection principles are at play in complex genotype to phenotype maps.

## ACKNOWLEDGEMENTS

I thank Ard Louis for useful discussions and for initially posing the question and David McCandlish for insightful comments on a previous version of the manuscript.

## APPENDIX

The Fokker-Planck equation describing the stochastic dynamics of the gene frequencies of multiple alleles is given by

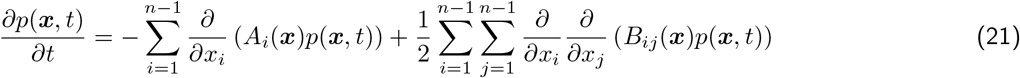

where *n* is the number of alleles, and the mean change in allele frequency (convective force) on the *i^th^* allele is

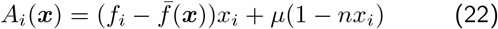

and the co-variance of the change in allele frequencies of the *i^th^* and *j^th^* allele (effective diffusion matrix) is

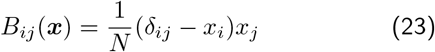

The equilibrium solution *p*^*^(***x**, t*) can be found bysettingthe flux ***J***(***x***) = 0, where the flux is defined by the continuity equation to be *∂_t_p*(***x**, t*) = −**∇ · J**:

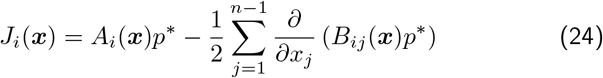

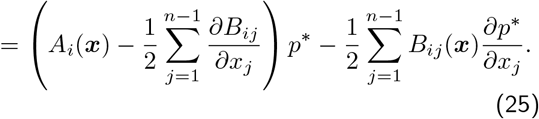

Multiplying through by the inverse 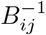 and summing it is simple to show that

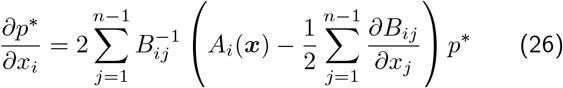

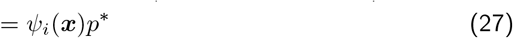

Now if the vector field is conservative, i.e. it is the gradient of a scalar function, ***ψ*** = **∇**Ψ(***x***), then the solution to Eqn.26 can be found using the standard integrating factor method:

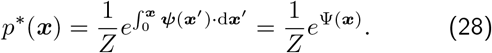

The vector field ***ψ*** is conservative if it is free from rotation, which in 3 dimensions or less means it is curl free. In higher dimensions an equivalent condition is:

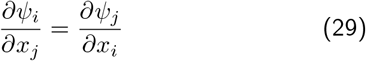

which we loosely refer to as the “curl-free” condition. We can evaluate *ψ_i_*(***x***), by using the fact that the inverse of *B_ij_* is given by^15^:

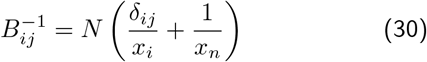

where 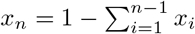 to give

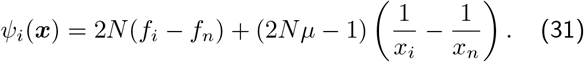

Evaluating the partial derivative wrt *x_j_* we find:

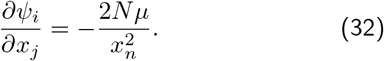

As this does not depend on *x_i_* or *x_j_* then this satisfies Eqn.29 and ***ψ*** is a curl-free vector field. Note that this is essentially a restatement of the fact that in order to find an equilibrium solution we are assuming detailed balance is obeyed. To find Ψ(***x***) we can integrate by inspection to get:

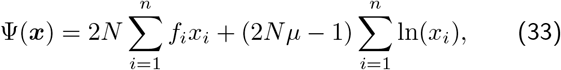

where note that the summations run to the index *i* = *n*. Plugging into Eqn.28, we find our final result

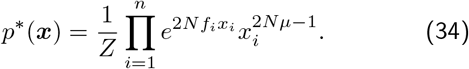

To the author’s knowledge this is the first time this derivation has appeared explicitly in the literature.

